# Single voxel autocorrelation reflects hippocampal function in temporal lobe epilepsy

**DOI:** 10.1101/2023.12.15.571916

**Authors:** Nichole R. Bouffard, Sam Audrain, Ali M. Golestani, Morgan D. Barense, Morris Moscovitch, Mary Pat McAndrews

## Abstract

We have previously shown that autocorrelation analyses of BOLD signal applied to single voxels in healthy controls can identify behaviorally-relevant gradients of temporal dynamics throughout the hippocampal long-axis. A question that remains is how changes in the brain’s functional and structural integrity affect single voxel autocorrelation. In this study we investigate how hippocampal autocorrelation is affected by hippocampal dysfunction by investigating a population of patients with unilateral temporal lobe epilepsy (TLE). Many patients with TLE have mesial temporal sclerosis (MTS), characterized by scarring and neuronal loss particularly in the anterior hippocampus. Here we compared patients with left and right TLE, some with and without MTS, to healthy controls. We applied our single voxel autocorrelation method and data-driven clustering approach to segment the hippocampus based on the autocorrelation of resting state fMRI. We found that patients with left TLE had longer intrinsic timescales (i.e., higher autocorrelation) compared to controls, particularly in the anterior-medial portion of the hippocampus. This was true for both the epileptogenic and non-epileptogenic hemispheres. We also evaluated the extent of cluster preservation (i.e., spatial overlap with controls) of patient autocorrelation clusters and the relationship to verbal and visuospatial memory. We found that patients with greater cluster preservation in the anterior-medial hippocampus had better memory performance. Surprisingly, we did not find any effect of MTS on single voxel autocorrelation, despite the structural changes associated with the condition. These results suggest that spatiotemporal dynamics of activity can be informative regarding the functional integrity of the hippocampus in TLE.

## 1. Introduction

Human functional MRI is a powerful neuroimaging tool that has enabled us to observe in-vivo timescales of brain activity. Recently our lab has developed an fMRI analytic technique that allows us to measure temporal dynamics in the BOLD signal over time in single voxels. This method, single voxel autocorrelation, is the lagged autocorrelation of individual voxels, which serves as a measure of how similar a voxel’s activity is to itself over time (Bouffard, Golestani et al., 2023; Coughlan et al., 2023). In healthy populations, single voxel autocorrelation is thought to reflect underlying neuronal timescales of activity, where high autocorrelation indicates longer timescales and low autocorrelation indicates short timescales (Gao et al., 2020; Watanabe et al., 2019; Wolff et al., 2022, Xie et al., 2023). Prior work from our lab identified gradients of single voxel autocorrelation within the hippocampus, along the anterior-posterior and medial-lateral axes, and showed that these gradients were related to behavior in a navigation task (Bouffard, Golestani et al., 2023). Gradients of single voxel autocorrelation, therefore, may be an important property of a healthy functioning brain and may reflect key aspects of cognitive processes in the hippocampus.

Disruption of hippocampal autocorrelation gradients could have significant consequences for cognition. Hippocampal functioning plays a central role in memory encoding, consolidation, and retrieval, and these processes depend not only on structural integrity, but also the coordination of interactions between the hippocampus and the broader memory network. Temporal lobe epilepsy (TLE), particularly cases with mesial temporal sclerosis (MTS), is a condition characterized by structural and functional abnormalities in the medial temporal lobes as well as cognitive deficits (Ives-Deliperi & Butler, 2021; McCormick et al., 2014). Studying this population, therefore, offers a unique opportunity to examine how altered hippocampal neural timescales relate to functional outcomes. In this study, we investigated patients with unilateral (left or right) TLE to determine how medial temporal lobe damage affects single voxel autocorrelation and how the spatial organization of autocorrelation relates to memory performance.

TLE is the most common type of drug-resistant epilepsy in adults, and is frequently associated with structural abnormalities in the hippocampus (Coras & Blümcke, 2015), including decreased volume and abnormal signal intensity on MRI, especially in the anterior hippocampus (Bernasconi et al., 2003; Jackson et al., 1990; Kuzniecky et al., 1997; Thom et al. 2012). These neuroimaging markers reflect MTS, characterized by hippocampal atrophy and gliosis present in approximately 50% of TLE patients (Bernasconi et al., 2003; Bernhardt et al., 2016; Kuzniecky et al., 1997; Thom et al., 2012). We hypothesized that such structural alterations may disrupt the temporal dynamics of hippocampal activity, reflected in changes in both single voxel autocorrelation and the organization of single voxel autocorrelation throughout the hippocampus relative to healthy controls. In addition to structural changes, unilateral TLE is also associated with lateralized memory impairments, where deficits in verbal memory are related to TLE in the left temporal lobe and deficits in visuospatial memory are related to TLE in the right temporal lobe (Kelley et al., 1998; Kim et al., 2003; St-Laurent et al., 2014). If single voxel autocorrelation indexes functional integrity, then abnormalities might be related to verbal and visuospatial memory performance in patients with TLE.

Other studies have examined neural signal dynamics in TLE using subdural EEG and have shown elevated temporal autocorrelation and reduced signal complexity in the epileptogenic hippocampus at rest (Monto et al., 2007; Parish et al., 2004) and during cognitive tasks (Protzner et al., 2010) relative to the healthy, contralateral hippocampus. Yet few studies have applied a single voxel autocorrelation approach to examine these intrinsic neural timescales in fMRI. A recent study by Xie et al., (2023) used a different autocorrelation approach and found a decrease in hippocampal autocorrelation in TLE, suggesting shorter timescales of activity in the hippocampus and more widespread areas of cortex bilaterally (Xie et al., 2023). These past studies, however, typically treat the hippocampus as a unitary structure, without considering the known spatial heterogeneity and its functional gradient organization (Vos de Wael et al., 2018). Our prior work suggests that the organization of hippocampal autocorrelation is meaningfully related to cognition (Bouffard, Golestani, et al., 2023). Thus, a key aim of the current study was to examine how location-dependent single voxel autocorrelation organization is affected by TLE and related to memory performance.

In healthy individuals, hippocampal single voxel autocorrelation follows a gradient from high-to-low along the anterior-posterior and medial-lateral axes (Bouffard, Golestani et al., 2023; Coughlan et al., 2023; Raut et al., 2020). Using data-driven clustering, these gradients can be segmented into three distinct clusters: an anterior-medial cluster of high autocorrelation voxels, an intermediate cluster, and a posterior-lateral cluster of low autocorrelation voxels (Bouffard, Golestani et al., 2023; Coughlan et al., 2023). Previous findings have shown that increases in autocorrelation in the anterior-medial cluster were predictive of increasing task difficulty during navigation, suggesting that both the magnitude and spatial location of autocorrelation matter for behavior. It remains unclear whether hippocampal damage in TLE disrupts the autocorrelation cluster organization and whether such disruption is functionally meaningful. In the current study, we evaluated changes in spatial organization of autocorrelation clusters by quantifying the spatial overlap of clusters between patients and controls and tested whether this overlap was related to memory performance.

Here we used resting-state fMRI to analyze hippocampal single voxel autocorrelation and autocorrelation clustering in patients with unilateral TLE and age-matched controls. We first tested whether the average single voxel autocorrelation and autocorrelation clusters differed between groups, with the prediction that the anterior clusters in patients, especially those with MTS, would be altered relative to controls.

Next, we correlated single voxel autocorrelation values with memory performance on neuropsychological tests. Finally, to determine whether the organization of single voxel autocorrelation is related to memory, we quantified cluster preservation (a measure of spatial overlap between patient and controls clusters) and tested its relationship to verbal and visuospatial memory. We predicted that patients with greater cluster preservation would have better memory performance, similar to other work in our lab showing that preservation of network functional connectivity is related to better memory (Barnett et al., 2019; Audrian et al., 2023).

By situating voxel-wise hippocampal temporal dynamics within a spatial and cognitive framework, this work aims to advance our understanding of how local timescales of neural activity relate to structural pathology and memory function in TLE. Discerning these relationships may provide complementary insights into functional integrity to our current suite of imaging tools in TLE.

## 2. Materials and Methods

### 2.1 Participants

Forty-nine adult patients with pharmacologically intractable unilateral TLE were recruited from the Epilepsy Clinic at Toronto Western Hospital. Twenty-three patients presented with left TLE (LTLE) (Female = 15, Mean Age = 37.43, SD = 12.46) and 26 presented with right TLE (RTLE) (Female = 9, Mean Age = 36.80, SD = 14.12).

Continuous recording of scalp EEG and video monitoring during an inpatient evaluation in our epilepsy monitoring unit were used to determine seizure focus. MTS was diagnosed based on designations by a neuroradiologist from patients’ T1- and T2-weighted anatomic scans. Fifteen patients with LTLE had MTS and 16 patients with RTLE had MTS. Two individuals (1 LTLE, 1 RTLE) did not have confirmed MTS but had hippocampal abnormalities. There was also one patient who had RTLE but MTS in the left hemisphere. Twenty-nine neurologically healthy control participants (Female = 16, Mean Age = 34, SD = 10.81) were recruited to serve as comparison for our patient sample. They were matched with the patient groups for sex and age. All controls gave prospective written informed consent. Prospective written informed consent was obtained from a subset of the patient group, while permission for retrospective analysis of clinical data (both neuropsychological and resting-state fMRI) was obtained from the University Health Network Ethics Board for a group of participants who were scanned prior to the current ethics protocol implementation.

A group-level summary of the clinical characteristics and demographics of the sample is presented in Table 1 and characteristics of each individual patient can be found in the Supplemental material (Table S1). The characteristics reported include laterality (left or right), sex, age of seizure onset, disease duration, Engel scores (taken at follow-up M = 3.64 years post-surgery), and MRI findings together with pathological confirmation, where available.

**Table 1.** Clinical characteristics and demographics of controls and patients.

| Characteristic | Controls | Left TLE | Right TLE | p |
| --- | --- | --- | --- | --- |
| n | 29 | 23 | 26 | – |
| Preoperative age, years (SD) | 34 (10.81) | 37.43 (12.46) | 37.04 (13.89) | 0.54 |
| Education, years (SD) | 17.9 (3.24) | 14.63 (2.65) | 14.56 (4.25) | < 0.001 |
| Sex, F/M | 16/13 | 15/8 | 9/17 | 0.09 |
| Language dominance, L/R/B/NA | – | 18/0/5/0 | 23/0/1/2 | 0.07 |
| Age at seizure onset, years (SD) | – | 17.23 (15.70) | 20.88 (12.61) | 0.37 |
| Disease duration, years (SD) | – | 20.21 (14.92) | 16.15 (14.23) | 0.34 |
| Had surgery, yes/no | – | 21/2 | 19/7 | 0.10 |
| Post-surgery outcome, seizure free/continued seizures, auras | – | 19/2 | 17/2 | 0.18 |
| MTS detected on MRI, L/R/Possible abnormality | – | 15/0/1 | 1/16/1 | 0.93 |
| MTS histopathologically confirmed, yes/no | – | 9/6 | 13/4 | 0.46 |

Designation of epileptogenic zones was determined based on electroencephalogram (EEG) recordings indicating exclusive temporal lobes onsets to seizures and interictal abnormalities. Some patients had extra MTL abnormalities (see Supplemental materials, Table S1). Hippocampal sclerosis was first determined based on radiological examination. The participants who underwent surgery also had their MTS status confirmed with histopathology. We conducted our primary analyses with patients that had MTS radiologically determined and then performed a secondary analysis with only the patients that had MTS confirmed with histopathology.

Language dominance for patients was determined with a clinical fMRI protocol. Patients were excluded from the current study if they had right hemisphere language dominance. There were two RTLE patients who had too much motion during their localizer scan, and language lateralization was uninterpretable in these two patients, but other clinical data was compatible with typical (left-hemisphere) dominance in these cases.

The central goal of our investigation was to determine if there was a relationship between hippocampal autocorrelation and verbal and visuospatial memory scores because these scores are representative of the material-specific memory loss typically found in left and right TLE cohorts (St-Laurent et al., 2014). These are the analyses that are presented in the paper. We also explored whether hippocampal autocorrelation might be related to other clinical characteristics particularly disease duration, but none were significant (see Supplemental materials, Page 16).

### 2.2 Neuropsychological Testing

A comprehensive neuropsychological battery was administered to patients that included assessment of intelligence, learning/memory, processing speed, and verbal and visuospatial functioning. Some patients elected to have surgery, and these patients completed the neuropsychological battery again approximately 1 year after surgery. The battery included the following measures: Wechsler Abbreviated Scale of Intelligence, Warrington recognition memory test for faces, Rey visual design learning test, spatial conditional associative learning test, Warrington recognition memory test for words, and Rey auditory verbal learning test. Summary memory scores were computed to capture a more reliable representation of core abilities than afforded by single test scores. We transformed the neuropsychological test scores into summary verbal and visuospatial scores using the previously estimated factor loadings from a principal components analysis following previous work from our group (see St-Laurent et al., 2014 for a full description of method). The verbal memory factor was based on loadings from correct responses on the Warrington recognition memory test for words (RMW), total recall (RAVLT-tot) over five study-test trials from the Rey auditory verbal learning test (RAVLT) and percent retention (RAVLT-ret) at 20-minute delay from the RAVLT. The visuospatial memory factor was primarily based on loadings from correct responses on the Warrington recognition memory test for faces (RMF), total recall across trials one through five on the Rey visual design learning test (RVDL), and number of trials to criterion for the conditional associative learning test (CAL). Past work from our group using this data reduction technique to generate the composite verbal and visuospatial measures has demonstrated that it is a powerful metric that can be reliably used to characterize the core abilities in TLE better than any one single neuropsychological test score (Audrain et al., 2018; Audrain et al., 2023; Barnett et al., 2015; Barnett et al., 2019; McCormick et al., 2014; St-Laurent et al., 2014). We examined the relationship between hippocampal autocorrelation and pre-surgery verbal and visuospatial memory scores as well as the relationship between autocorrelation and the change in memory scores (pre- to post-surgery), in patients that had surgery which at our center was a standard anterior temporal lobe resection including the amygdala and hippocampus.

### 2.3 MRI Acquisition

A high-resolution 3D anatomical scan was collected on a 3T Signa MR system (GE Medical Systems, Milwaukee, WI, USA) for each participant (T1-weighted sequence, FOV 220mm, 146 slices, flip angle = 12 degrees, 256×256 matrix, resulting in voxel size of 0.86 x 0.86 x 1.0). Six-minute resting state fMRI (T2∗-weighted) scans were acquired with an echo-planar pulse imaging (EPI) sequence (FOV 240mm, 28–32 slices depending on head size, TR = 2,000ms, TE = 25ms, 64 x 64 matrix, 3.75 x 3.75 x 5mm voxels). During resting state scans, participants were instructed to lie still, and “not to think about anything in particular,” with their eyes closed.

### 2.4 Functional MRI Pre-processing

Preprocessing was performed using SPM8 (http://www.fil.ion.ucl.ac.uk/spm/software/spm8), a toolbox running in MATLAB 2018a (Mathworks). Anatomical and functional images were reoriented so that the origin falls on the anterior commissure. The functional images were then corrected for slice-timing, realigned and unwarped, and then co-registered to the anatomical image. Functional and anatomical images for each participant were normalized into standard Montreal Neurological Institution (MNI) space, segmented into gray matter, white matter and cerebral spinal fluid (CSF), and were resampled to 2 mm isotropic voxels following a directed normalization procedure (Calhoun et al., 2017; Nieto-Castanon, 2022) using SPM unified segmentation and normalization algorithm (Ashburner, 2005; Ashburner & Friston, 2005). We resampled the data to enable alignment to a common MNI space. The single voxel autocorrelation method involves analysis of individual voxels therefore no spatial smoothing was performed. Using the Conn toolbox (Whitfield-Gabrieli & Nieto-Castanon, 2012), a CompCor (Behzadi et al., 2007) was used to exclude measures of physiological noise by regressing out the top five components of a principal components analysis from eroded white matter and CSF masks produced from the SPM8 segmentation. Motion scrubbing using the ART function was also applied to reduce artefacts due to motion. Any timepoint that was 3 or more standard deviations from the mean global brain activation, more than 1mm of linear motion, or more than 0.05 radians rotation was regressed from the data. Low-pass temporal filtering was applied before computing the single voxel autocorrelation to exclude high (>0.09Hz) frequency fluctuations. The filtered and corrected images were used for subsequent analyses.

### 2.5 Single voxel autocorrelation method

#### 2.5.1 Computing single voxel autocorrelation

We used the single voxel autocorrelation method described by Bouffard, Golestani et al. (2023) and describe the method briefly here. Bilateral hippocampal masks were from the Brainnetome Atlas (Fan et al., 2016) (BNA), which had 1161 voxels in the left hemisphere and 1098 voxels in the right hemisphere. For each voxel, unbiased autocorrelation (as implemented in MATLAB xcorr cross-correlation function) was calculated. Specifically, the timecourse of a single voxel’s activity was correlated with itself shifted by a temporal lag, the length of 1 TR (TR = 2000 ms). We repeated this process, shifting the timecourse forward by 1 lag (2000 ms) and correlating it with the original, non-shifted timecourse until a maximum temporal shift of 4 seconds was reached. The autocorrelation of the BOLD signal in the gray matter drops off after approximately 4000 ms (or 4 second) (i.e., it is not distinguishable from the autocorrelation of other noise) (Bollmann et al., 2018; Watanabe et al., 2019). In the current dataset, the TR was 2 seconds, therefore maximum lag shift possible (without exceeding 4 seconds) is 2 lags. The non-shifted timecourse was correlated with lag 1 (length of 1 TR) and with lag 2 (length of 2 TRs) (Figure 1A). The autocorrelation (AC) computed for each lag was stored in a vector. The autocorrelation vector contained 2 values (one single voxel autocorrelation value for each lag). This approach resulted in an autocorrelation vector for each voxel (Figure 1). The vectors were then used to compute autocorrelation clusters in the next step.

**Figure 1.**
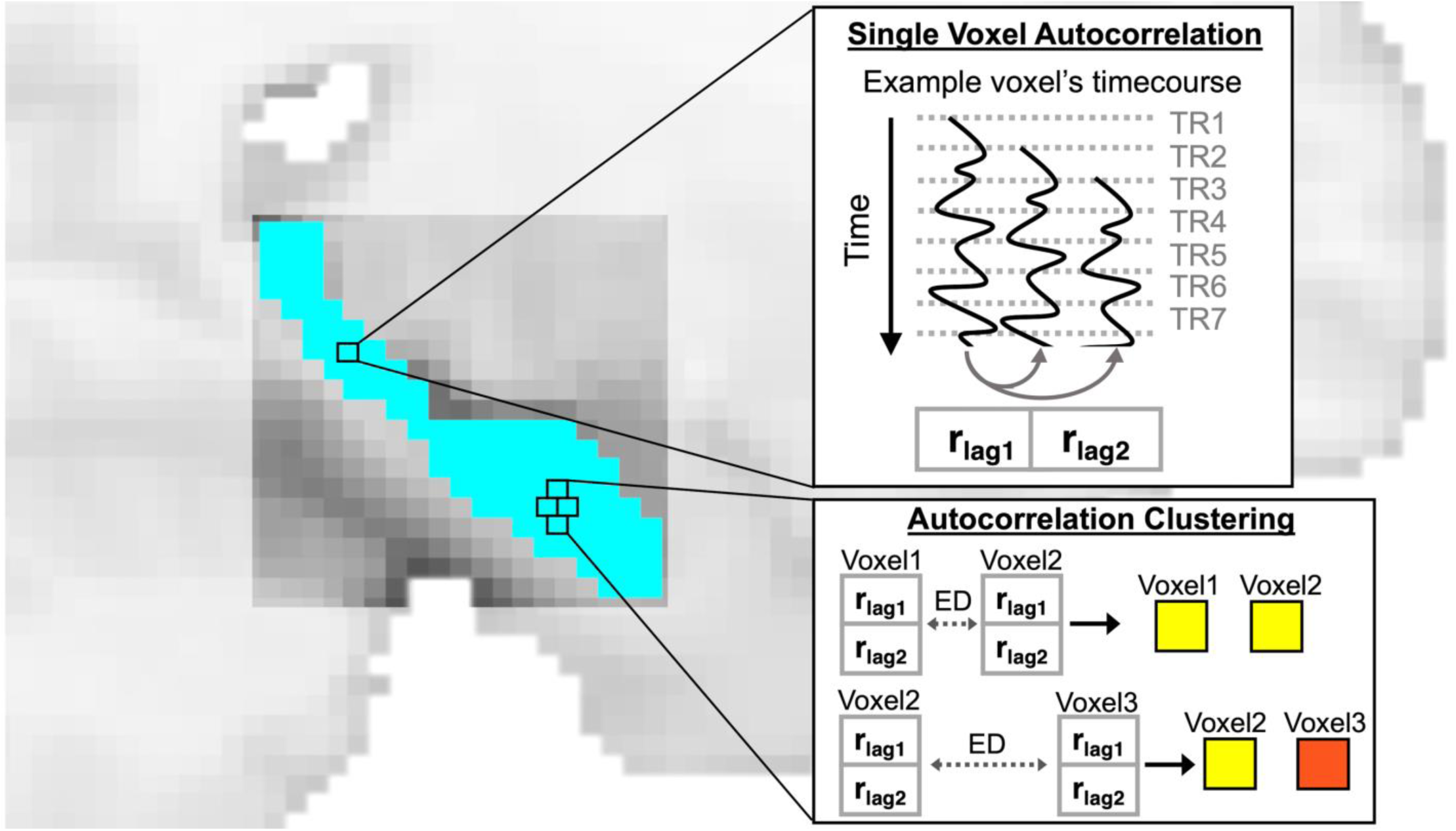
Single voxel autocorrelation method. For each voxel in the hippocampus, the time course of BOLD activity was successively temporally shifted by 1 TR and correlated with itself. This was repeated for a total shift of 4 seconds (i.e., 2 lags). This resulted in a vector of single voxel autocorrelation values, with each value corresponding to a different lagged correlation. **Autocorrelation clustering.** The single voxel autocorrelation vectors for every voxel were used to compute the autocorrelation clusters, where voxels were clustered based on the Euclidean distance (ED) of their single voxel autocorrelation vectors. Every voxel was assigned to a cluster and the total number of clusters was determined by the modularity optimization algorithm (i.e., number of clusters was not predetermined). Yellow and red voxels represent voxels that were assigned to the same (voxel 1 and 2) or different clusters (voxel 1 and 3). Clustering was performed at the individual participant level, and a separate cluster map was created for each individual and each hemisphere. Group-level cluster maps were created by averaging across individuals.

#### 2.5.2 Autocorrelation Clustering

For the next step, the Euclidean distance between the single voxel autocorrelation vectors of each voxel pair in each mask was calculated to create a distance matrix. The distance matrix was first normalized (i.e., divided by the maximum value) and then subtracted from 1 to generate a similarity matrix ranging from 0 to 1.

This similarity matrix was used to generate hippocampal clusters using the modularity optimization algorithm proposed by Blondel et al., (2008) and Wickramaarachchi et al., (2014). Unlike most other clustering methods, modularity optimization does not require a predesignated number of clusters and instead estimates the optimum number of clusters from data. An autocorrelation cluster map for left and right hippocampus was created for each individual. In addition to clustering at the individual level, group-level clusters were derived by averaging the similarity matrices of all participants. Group-level autocorrelation cluster maps were created for the Control, LTLE and RTLE groups (Figure 2).

**Figure 2.**
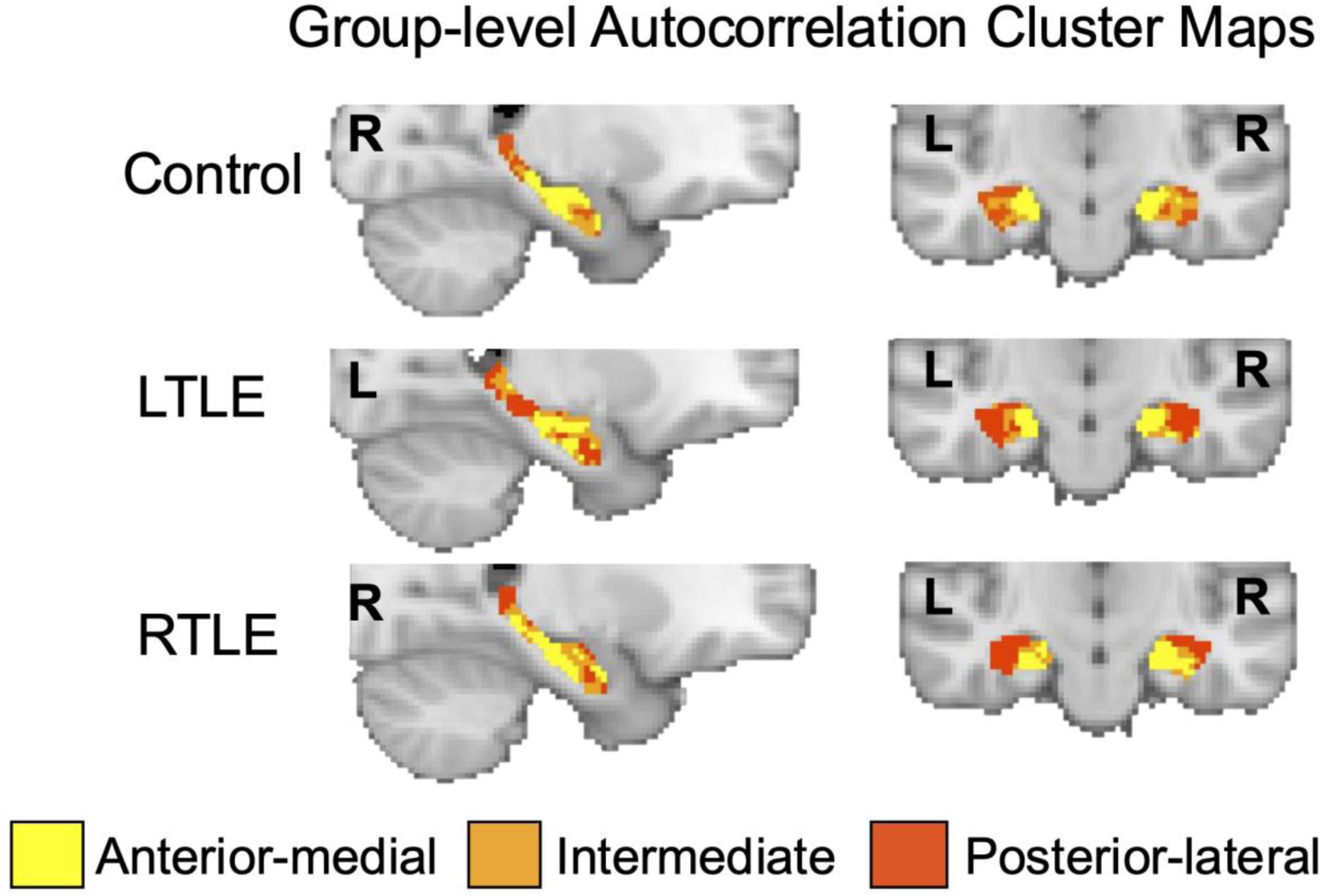
Group-level autocorrelation clusters. Hippocampal voxels were clustered based on the similarity (Euclidean distance) of single voxel autocorrelation vectors, resulting in one cluster in the anterior-medial hippocampus (yellow) associated with high autocorrelation values, one in the posterior-lateral hippocampus (red) associated with low autocorrelation values, and one cluster in the intermediate hippocampus (orange) in between the anterior-medial and posterior-lateral clusters.

We validated the cluster maps in two ways. First, to assess whether the spatial organization of clusters was due solely to spatial autocorrelation or artifacts associated with low-resolution data, we implemented a spin-based permutation analysis (Burt et al., 2020) using the BrainSpace toolbox (Vos de Wael et al., 2020), where data from the Control group was used to generate surrogate datasets that preserved the spatial autocorrelation of the data, while removing spatial specificity. We applied the autocorrelation clustering to the surrogate data which then allowed us to create a null distribution of spatial overlap for each cluster. This distribution was used to determine the threshold for what values constitute “high” spatial overlap and spatial replication of clusters. This analysis demonstrated that the spatial organization of cluster maps derived from patient data are not solely due to artifacts (see Supplemental materials Figure S1).

Next, we computed reliability of autocorrelation clusters by computing the overlap of clusters among individuals within each group (Control, LTLE, RTLE). By comparing the spatial organization of clusters of individuals within the same group, we determined that the anterior-medial cluster and posterior-lateral cluster were the most reliable clusters, whereas the intermediate cluster was more variable across individuals and therefore less reliable. See Supplemental Materials for the detailed reliability analysis and Figure S2 for the reliability data of each cluster.

### 2.6 Cluster preservation: Overlap of individual autocorrelation clusters with group-level Control clusters

To compute cluster preservation, or the overlap of individual patient clusters with the group-averaged Control clusters, we measured the spatial overlap (Jaccard coefficient) between each patient’s autocorrelation cluster map with the group-level average autocorrelation cluster map from the Control group. The Jaccard coefficient of regions A and B is defined as:

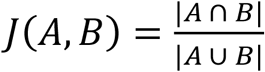

Where |*A* ∩ *B*| is the number of common voxels in both A and B (intersection) and |*A* ∪ *B*| is the number of voxels in A and B combined (union). We calculated the Jaccard coefficient between each individual’s autocorrelation cluster map and the group-level autocorrelation cluster map from Controls. We computed the Jaccard coefficient for each autocorrelation cluster separately.

High spatial overlap (i.e., large Jaccard coefficients) suggests greater spatial similarity/preservation of the spatial distribution of autocorrelation clusters relative to Controls. Low spatial overlap (i.e., small Jaccard coefficients) would suggest less spatial similarity/preservation of spatial distribution of autocorrelation clusters relative to Controls. For example, consider a voxel from the healthy control group that had high single voxel autocorrelation and was assigned to a specific cluster. In a patient, if that same voxel had a decrease in single voxel autocorrelation due to changes in the underlying dynamics, that voxel would now be assigned to a different cluster. This assignment would lead to changes in how the autocorrelation clusters are organized throughout the hippocampus of patients compared to controls.

To determine whether cluster preservation was related to memory, we correlated cluster preservation and patient neuropsychological test summary scores for verbal and visuospatial memory. Left TLE is associated with deficits in verbal memory and right TLE is associated with deficits in visuospatial memory (Kim et al., 2003; St-Laurent et al., 2014). Therefore, we specifically correlated verbal memory scores with cluster preservation from LTLE patients and visuospatial memory scores with cluster preservation from RTLE patients.

## 3. Results

### 3.1 Single voxel autocorrelation clusters

We found that nearly every individual within the Control, LTLE, and RTLE groups had three hippocampal autocorrelation clusters in both the left and right hemispheres (Figure 2). There was one exception, a Control participant who only had two autocorrelation clusters in the right hemisphere, and we excluded this participant from further analyses. Consistent with prior work, we found one cluster in the posterior-lateral hippocampus, one cluster in the anterior-medial hippocampus, and an intermediate cluster (Figure 2). We also computed the size of each cluster (i.e., number of voxels assigned to each cluster). There were no significant differences in cluster size between Control, LTLE, and RTLE groups (See Supplemental Materials Table S2 for list of cluster sizes).

For the following analyses, we computed the average single voxel autocorrelation in each of the three clusters per participant. We ran a linear mixed effects model with cluster (anterior-medial, intermediate, posterior-lateral), group (LTLE (N=23), RTLE (N=26), Control (N=28)), and hemisphere (left, right) as predictors, including participant as the random intercept in the random effects term. We found a main effect of cluster (F(2,370) = 378.25, p <0.001) and a main effect of group (F(2,74)=5.15, p <.01). There was no effect of hemisphere (F(1,370)= 0.78, p = 0.37). For the post hoc analysis of the main effects, we used the Kenward-Roger method to calculate the degrees of freedom and the Tukey method to correct for multiple comparisons. Post-hoc investigations of the main effect of cluster revealed that the anterior-medial cluster had the highest single voxel autocorrelation, followed by the intermediate cluster, and lastly the posterior-lateral cluster, which had the lowest single voxel autocorrelation (anterior-medial > intermediate: t(370)= 20.95, p < 0.001; anterior-medial > posterior-lateral: t(370) = 25.91, p < 0.001; intermediate > posterior-lateral: t(370) = 4.96, p < 0.001) (Figure 3A). See Supplemental Materials Table S3 for average autocorrelation values per cluster. Post-hoc analysis of the main effect of group revealed that individuals with LTLE had higher single voxel autocorrelation values than those with RTLE or Controls (LTLE > RTLE: t(74) = 2.76, p < 0.05; LTLE > Controls: t(74) = 2.87, p < 0.05). There was no difference in single voxel autocorrelation between the RTLE and Control groups (t(74) = 0.06, p=0.99).

**Figure 3.**
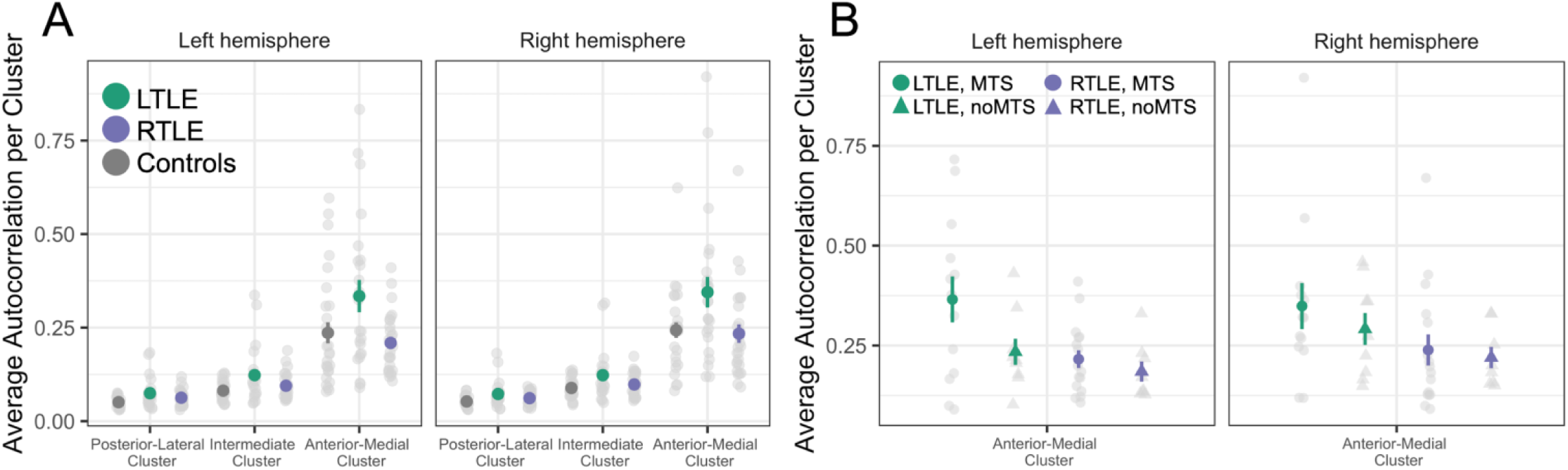
A) Average autocorrelation per cluster by group. The single voxel autocorrelation was averaged across voxels within each of the autocorrelation clusters from individuals in each of the three groups: Controls (gray), LTLE (green), and RTLE (purple). **B) Average autocorrelation in the anterior-medial hippocampal cluster by group and MTS status.** The average single voxel autocorrelation in just the anterior-medial cluster for LTLE and RTLE patients. Individuals are grouped by whether they had confirmed hippocampal mesial temporal sclerosis (MTS), resulting in four groups: LTLE with MTS (green circle), LTLE with noMTS (green triangle), RTLE with MTS (purple circle), and RTLE with noMTS (purple triangle).

We also found a significant cluster x group interaction (F(4,370)=8.95, p < 0.001). Post-hoc analysis revealed that this interaction was driven by higher single voxel autocorrelation in the anterior-medial cluster in LTLE patients compared to RTLE and Controls (LTLE > RTLE: t(134) = 5.35, p < 0.001; LTLE > Controls: t(134) = 4.61, p < 0.001) (Figure 3A). Notably, this was true for both the epileptogenic (left) and non-epileptogenic (right) hippocampi.

These findings are consistent with our prior work in healthy individuals which has shown that clusters located in the anterior-medial hippocampus are associated with higher autocorrelation values compared to clusters in the posterior-lateral hippocampus. We predicted that both LTLE and RTLE groups would differ from Control and that within patients there would be a difference between affected and healthy hippocampi. In contrast, only the LTLE group had higher single voxel autocorrelation than controls, and that was seen in both left and right anterior-medial clusters.

In past work, the intermediate cluster has been shown to have low reliability within participants and includes significantly fewer voxels compared to the anterior-medial and posterior-lateral clusters (Bouffard, Golestani et al., 2023; Coughlan et al., 2023). Here we again replicate this finding and found that the intermediate cluster had low reliability (See Supplemental Materials Figure S2) and was the smallest cluster (See Supplemental Materials Table S2 for list of cluster sizes). Therefore, for the rest of the analyses in the paper we chose to exclude it and only focus on the anterior-medial and posterior-lateral clusters.

To test empirically whether the autocorrelation clusters maps were a replication of past work, we compared group-level clusters maps (Controls, LTLE, RTLE) to the group-level cluster maps from a resting state dataset (a subset of the Human Connectome Project Retest dataset (N = 44 participants), see Bouffard, Golestani, et al., 2023 for more details). We focused our analyses on the anterior-medial and posterior-lateral clusters (our main clusters of interest) and examined the left and right hemisphere separately. Clusters were considered to be replicated if they have high overlap, or Jaccard coefficients greater than 0.35 in the anterior-medial cluster and greater than 0.42 in the posterior-lateral cluster. These were determined to be the 95th percentile based on the null distribution of spatial overlap (see Supplemental materials Figure S1).

In Controls, we found a replication of our prior clusters in both left and right hemispheres (Table 2). In LTLE and RTLE patients, we also found replication of all clusters in both hemispheres (Table 2). Interestingly, the clusters with the numerically lowest coefficients were the anterior-medial cluster in the epileptogenic hippocampus. The anterior-medial cluster in the left hemisphere of LTLE patients had a Jaccard coefficient equal to 0.39 and the anterior-medial cluster in the right hemisphere of RTLE patients had a Jaccard coefficient equal to 0.37. While these values still exceed our 95th percentile cut-off of 0.35, this decrease in overlap might suggest that spatial location of the anterior-medial cluster of the epileptogenic hippocampus is disrupted in TLE.

**Table 2.** Spatial overlap of autocorrelation clusters (Jaccard coefficients) relative to prior dataset from Bouffard, Golestani, et al., *Cerebral Cortex*, 2023.

|  | **Left Hippocampus** |  | **Right Hippocampus** |  |
| --- | --- | --- | --- | --- |
|  | **Anterior-medial** | **Posterior-lateral** | **Anterior-medial** | **Posterior-lateral** |
| Controls | 0.44 | 0.49 | 0.41 | 0.51 |
| Left TLE | 0.39 | 0.54 | 0.48 | 0.58 |
| Right TLE | 0.49 | 0.51 | 0.37 | 0.56 |

#### 3.1.1. Effect of Mesial Temporal Sclerosis (MTS) on single voxel autocorrelation

We predicted that hippocampal autocorrelation in patients with MTS would be different compared to patients without MTS in the affected hippocampus. Given the loss of neurons and associated gliosis is typically located in the anterior hippocampus (based on radiology examinations), we restricted our analyses to the cluster located in the anterior-medial hippocampus, as we predicted it would be more affected than the other clusters. We ran a linear mixed effects model on the average single voxel autocorrelation in the anterior-medial cluster for the LTLE and RTLE groups. We included MTS (MTS, noMTS), group (LTLE, RTLE), and hemisphere (left, right) as predictors. Note, there were 2 people (1 LTLE, 1 RTLE) who did not have confirmed MTS but had possible hippocampal abnormalities. Because we were not sure whether these participants had MTS, we excluded them from this analysis. There was also one participant who had RTLE but MTS in the left hemisphere, so we also excluded that individual from this analysis. With these exclusions, the number of participants included in the following analyses was LTLE (N=22) and RTLE (N=24).

As above, we found a significant effect of group (F(1,42) = 4.48, p <0.05), with the LTLE group demonstrating higher single voxel autocorrelation compared to the RTLE (t(42) = 2.11, p <0.05) (Figure 3B). Critically, we did not find an effect of MTS (F(1,42) = 1.69, p = 0.201) or any other significant effects or interactions. Nonetheless, individuals with confirmed MTS in the LTLE group had numerically higher single voxel autocorrelation in the affected hemisphere (left) than individuals without MTS. Given our predictions that MTS would affect single voxel autocorrelation, we tested this relationship directly with an independent samples t-test and did not find a significant difference in single voxel autocorrelation in the MTS group compared to those without MTS (t(18.99) = 1.57, p = 0.132). We also conducted a follow-up analysis in which we re-ran the previous model but only included patients who had MTS that was confirmed with histopathology (LTLE N = 9, RTLE N = 13). We found no significant effect of MTS (F(1, 33) = 2.00, p= 0.16) and no significant interactions.

We did not find a significant effect of MTS when we examined the average single voxel autocorrelation within the autocorrelation clusters. This finding is surprising, given that hippocampal atrophy is a defining characteristic of MTS. We investigated this further by examining whether hippocampal volume, in place of our binary variable for MTS, would reveal a relationship with autocorrelation. We did not find any significant effects of hippocampal volume. The full details of this analysis are described in the Supplemental materials, (Figure S3). Additionally, we hypothesized that the lack of effect of MTS could be due to low statistical power, therefore we conducted a flipped brain analysis where we normalized both groups to the control group and then used a grouping variable of “epileptogenic hemisphere” and “healthy hemisphere” instead of left and right hemisphere (Liu et al., 2015). We again did not find significant effects of epileptogenic vs. healthy hemisphere on the average autocorrelation. Together the results of these two supplemental analyses provide further evidence that single voxel autocorrelation is not directly related to hippocampal structural integrity. The full details of this analysis are described in the Supplemental materials, Page 17-18.

Lastly, we hypothesized that the clustering step might be obscuring fine-grained differences in autocorrelation between patients and controls and between patients with and without MTS. We conducted a voxel-level analysis in which we used the single voxel autocorrelation (lag 1) of each voxel as the dependent variable in the linear mixed effects models. Using the voxel-level approach, we found results similar to those reported above, and notably did not find significant differences in autocorrelation due to MTS. The detailed analysis is included in the Supplementary materials (Page 14-15).

#### 3.1.2. Relationship of single voxel autocorrelation to memory

To determine whether autocorrelation was related to memory we calculated the Pearson’s correlation between the average single voxel autocorrelation in each of the clusters with pre-surgery verbal memory and visuospatial memory scores. We did this for each hemisphere separately as the well-known lateralization of verbal memory in the left temporal lobe and nonverbal memory in the right temporal lobe led us to specific predictions (Kelley et al., 1998; Kim et al., 2003; St-Laurent et al., 2014). We predicted that single voxel autocorrelation in the epileptogenic hemisphere of patients would be correlated with the corresponding dominant memory process attributed to that hemisphere (e.g., verbal memory in LTLE and visuospatial memory in RTLE). Note, one patient from the RTLE group had to be excluded from this analysis because we did not have all the relevant neuropsychological test measures for PCA. With this exclusion, the number of participants included in this analysis was LTLE (N=23) and RTLE (N=25). We did not find any significant correlations between average single voxel autocorrelation and verbal memory in LTLE or visuospatial memory in RTLE, in either cluster (range of Rs: 0.07 - 0.34). See Supplemental Materials Table S4 for all R and p values (corrected for multiple comparisons using FDR).

For the TLE patients who elected to have surgery, the neuropsychological battery was administered again post-surgery and verbal and visuospatial memory scores were computed. Although we did not observe a relationship between autocorrelation and pre-surgery memory scores, we investigated whether autocorrelation might be related to changes in memory scores from pre- to post-surgery. We first computed the change in verbal memory scores (pre-minus post-surgery) for the LTLE group and the change in visuospatial memory scores for the RTLE group. We then correlated the autocorrelation in the anterior-medial hippocampus with the change in memory scores. There were no significant correlations that survived multiple comparisons correction. The details and results of this analysis are described in the Supplemental materials Figure S3.

### 3.2 Cluster preservation

We predicted that the spatial organization of autocorrelation throughout the hippocampus would be disrupted in cases of TLE. To test this prediction, we measured cluster preservation by comparing the spatial organization of individual patients’ autocorrelation clusters to the spatial organization of group averaged Control clusters. We calculated the Jaccard coefficient, or the spatial overlap, between each patient’s autocorrelation cluster map and compared it to the group-level autocorrelation cluster map for the Control group. The group-level autocorrelation cluster map was computed from Control participants (N=28) who all had three hippocampal clusters. A high Jaccard coefficient suggests that the spatial organization of autocorrelation clusters for an individual is comparable to the Control group and has high cluster preservation, whereas a low coefficient suggests the spatial organization has shifted compared to the normative group. Jaccard coefficients were considered high if they were greater than 0.35 in the anterior-medial cluster and greater than 0.42 in the posterior-lateral cluster. These values were determined based on the null distribution of spatial overlap (see Supplemental materials Figure S1). Also see Supplemental Materials Table S5 for the Jaccard coefficients values for each autocorrelation cluster.

We conducted a linear mixed effects model on the Jaccard coefficients and included group (LTLE (N=23), RTLE (N=26), cluster (anterior-medial, posterior-lateral), and hemisphere (Left, Right) as the fixed effects predictors. We found a main effect of cluster (F(1,141) = 34.36, p <0.001) and a main effect of hemisphere (F(1,141) = 42.59, p <0.001). We did not find a main effect of group (p = 0.594). We also found a significant cluster x hemisphere interaction (F(1,141) = 8.87, p <0.01).

Post-hoc analysis of the main effect of cluster revealed that the anterior-medial cluster had lower cluster preservation, and therefore a greater deviation from the Control group, compared to the posterior-lateral cluster (t(141) = 5.86, p < 0.001). Post hoc analysis of the main effect of hemisphere revealed that the right hemisphere had greater cluster preservation than the left (t(141) = 6.52, p < 001). We did not find any differences in cluster preservation between LTLE and RTLE. Analysis of the cluster x hemisphere interaction revealed that the left anterior-medial cluster had lower cluster preservation compared to the right anterior-medial cluster (t(141) = 6.72, p <.001) and compared to both left (t(141) = 6.25, p <.001) and right (t(141) = 8.75, p <.001) posterior-lateral clusters. For the post hoc analysis of the main effects, we used the Kenward-Roger method to calculate the degrees of freedom and the Tukey method to correct for multiple comparisons.

#### 3.2.1. Effect of MTS on cluster preservation

We predicted that patients with MTS would have decreased cluster preservation compared to those without MTS due to the structural changes caused by MTS. Because MTS typically affects the anterior hippocampus, we had specific predictions that the anterior-medial hippocampus would demonstrate more of a change in cluster preservation. To test this, we ran a linear mixed effects model on the Jaccard coefficients just in the anterior-medial cluster. We included MTS (based off radiology examinations) (MTS, noMTS), group (LTLE, RTLE), and hemisphere (left, right) as predictors. We again excluded the two people with possible abnormalities from this analysis and the one RTLE participant with left MTS. With these exclusions, the number of participants included in the following analyses was LTLE (N=22) and RTLE (N=24). We did not find a significant effect of MTS (F(1,42) = 0.75, p = 0.391), nor did we find that MTS interacted with the other variables in the model. We also conducted a follow-up analysis in which we re-ran the previous model but only included patients who had MTS that was confirmed with histopathology (LTLE N = 9, RTLE N = 13). We found no significant effect of MTS (F(1, 33) = 0.49, p =0.490) and no significant interactions. This result suggests that there was no difference in spatial organization of autocorrelation clusters between patients with MTS and without MTS.

#### 3.2.2. Relationship of cluster preservation to memory

To understand whether the cluster preservation of autocorrelation clusters was related to memory, we computed the Pearson’s correlation between cluster preservation and material-specific, lateralized pre-surgery memory scores (i.e., verbal memory scores for LTLE and visuospatial memory scores for RTLE). We focused our analyses on the anterior-medial cluster as that had shown the greatest deviation from the Control group. As above, one patient from the RTLE group had to be excluded from this analysis because we did not have their neuropsychology test scores. With this exclusion, the number of participants included in the following analyses was LTLE (N=23) and RTLE (N=25). We found that for LTLE individuals, better verbal memory scores were significantly correlated with greater cluster preservation in the anterior-medial cluster of the left hemisphere (R = 0.46, p < 0.05, p_corrected_ = 0.05), and with multiple comparisons correction (FDR) the p = 0.05 (Figure 4). We did not find a significant correlation in the right hemisphere of the LTLE patients (p = 0.64). We found greater cluster preservation of the anterior-medial cluster in the right hippocampus for the RTLE group was correlated with better visuospatial memory (R=0.44, p < 0.05, p_corrected_ = 0.05), and with multiple comparisons correction (FDR) the p = 0.05 (Figure 4). We did not find a significant correlation in the left hemisphere of the RTLE group (p = 0.88). Thus, there was a positive association between cluster preservation and material-specific memory performance in the epileptogenic hippocampus of both LTLE and RTLE patients.

**Figure 4.**
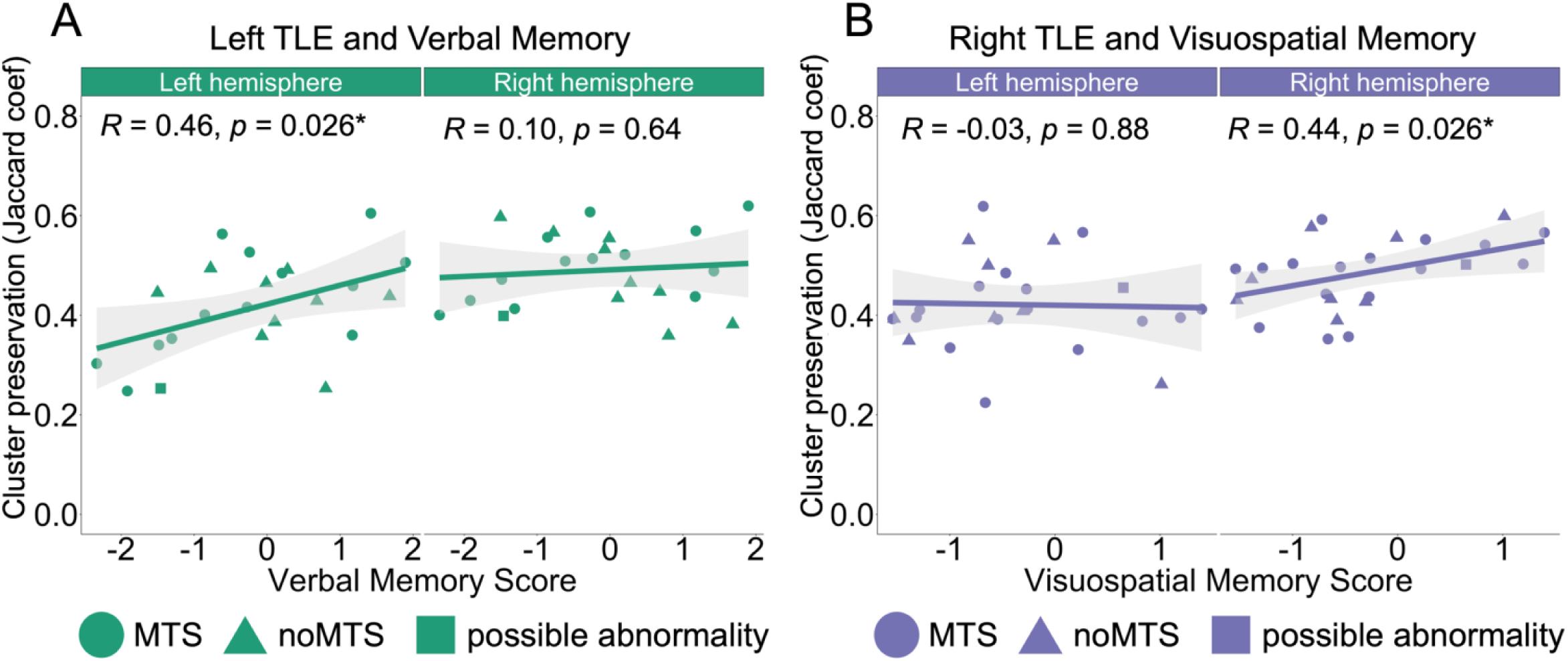
Cluster preservation in the anterior-medial cluster is correlated with memory. **A.** Verbal memory score correlated with cluster preservation (i.e., Jaccard coefficient) of the anterior-medial cluster in the left hemisphere and right hemisphere of LTLE individuals. The x-axis indicates verbal memory scores, where higher scores represent better performance. The y-axis indicates cluster preservation, where higher values represent greater similarity to Controls. LTLE patients with MTS (green circle), LTLE patients with noMTS (green triangle), and LTLE patients with possible hippocampal abnormality (green square). **B.** Visuospatial memory score correlated with cluster preservation of the anterior-medial cluster in the left and right hemisphere of individuals with RTLE. The x-axis indicates visuospatial memory scores, where higher scores represent better performance. The y-axis indicates cluster preservation, where higher values represent greater similarity to Controls. RTLE patients with MTS (purple circle), RTLE patients with noMTS (purple triangle), and RTLE patients with possible hippocampal abnormality (purple square).

We also investigated whether cluster preservation was related to changes in memory scores from pre- to post-surgery. We correlated the cluster preservation in the anterior-medial hippocampus with the change in verbal and visuospatial memory scores. There were no significant correlations that survived multiple comparisons correction. The details and results of this analysis are described in the Supplemental materials Figure S3.

## 4. Discussion

The aim of the current study was to understand how hippocampal single voxel autocorrelation is organized in individuals with unilateral TLE, and whether this measure captures structural pathology and memory function in this population. Here, we replicated our past work (Bouffard, Golestani et al., 2023; Coughlan et al., 2023) and found that all groups demonstrated a pattern of high autocorrelation values clustered in the anterior-medial hippocampus and low autocorrelation values clustered in the posterior-lateral. There were two major findings in the TLE groups, both selective to the anterior-medial cluster. First, those with left TLE showed higher single voxel autocorrelation, suggesting longer neural timescales, in comparison to other groups.

Second, the preservation of spatial organization of single voxel autocorrelation clusters was correlated with material-specific memory performance in both left and right TLE patients. Contrary to our predictions, we did not find significant differences in single voxel autocorrelation values or cluster preservation according to epileptogenic versus healthy hemispheres or as a function of radiologically-evident MTS. These findings suggest that single voxel autocorrelation as an index of hippocampal dynamics may be a more direct measure of functional rather than structural integrity, at least with regards to individuals with TLE.

The first main finding was that patients with LTLE had greater single voxel autocorrelation in the anterior-medial hippocampus, indicating longer intrinsic timescales of activity, compared to controls and to patients with RTLE. This finding is consistent with prior work that found slower decay rates of autocorrelation (i.e., greater autocorrelation) in the ipsilateral anterior hippocampus of TLE patients compared to controls (Nedic et al., 2015). Another recent study, however, examined autocorrelation in the hippocampus of patients with TLE and found a different result. Xie et al. (2023) found reduced autocorrelation in the hippocampus bilaterally, suggesting shorter timescales across the whole hippocampus in patients with TLE, which is the opposite of our findings and the findings in Nedic et al. There are methodological differences that could potentially account for the differences in results across studies. For example, while Nedic et al. examined the rate of decay of the autocorrelation function, Xie et al. computed intrinsic neural timescales, a measure of the positive area under the autocorrelation function curve. Both of which differ from our method, single voxel autocorrelation that applies a constant temporal window, or 2 temporal lags, resulting in a measure of the intrinsic timescale of activity over a fixed window of time. Differences in the calculation of these measures could potentially lead to mixed results. As there are relatively few studies examining region-specific intrinsic neural timescales and how they are affected in cases of TLE, future work is needed to determine which signal dynamic profiles most accurately represent this population.

The second main finding was that spatial preservation of the autocorrelation clusters was predictive of memory performance. By examining cluster preservation, we were able to combine both the temporal and spatial components of autocorrelation and examine how a divergence from the norm influences memory. This finding suggests that in patients, changes in the underlying signal dynamics were reflected in changes of the single voxel autocorrelation value, which led to different cluster assignment of hippocampal voxels in patients compared to controls. This difference in cluster assignment resulted in subtle changes in how the autocorrelation clusters were organized throughout the hippocampus of patients which we were able to quantify with our cluster preservation measure. Here, we also showed that the more ‘normative’ the organization of autocorrelation values, particularly in the anterior-medial hippocampus, the better the memory performance in a material-specific, lateralized manner.

In previous work correlating the temporal dynamics of the BOLD signal with behavior, task-dependent changes in signal dynamics have been associated with better cognitive performance within subjects (Armbruster-Genç et al., 2016; Bouffard, Golestani et al., 2023) and between subjects (Protzner et al., 2013). We had predicted that our measure, single voxel autocorrelation, could be an indicator of this aspect of functional integrity and thus would be related to patient’s memory performance. This prediction was not borne out for single voxel autocorrelation values and offline memory measures. One important caveat is that we were examining single voxel autocorrelation during resting state fMRI, whereas the previous studies relating signal dynamics and behavior used task fMRI (Armbruster-Genç et al., 2016; Bouffard, Golestani et al., 2023; Garrett et al., 2011, 2014; Protzner et al., 2013). It is plausible that single voxel autocorrelation measures neural variability related to real-time cognitive processes, as seen in our past work where autocorrelation changes were related to navigation task difficulty (Bouffard, Golestani et al., 2023; Brunec, Bellana et al., 2018), but that there is a weaker relationship between single voxel autocorrelation during rest and more trait-like evaluations of behavior. Thus, chronically longer intrinsic timescales may not be sufficient to explain variations in complex operations underlying offline performance on clinical memory measures.

Given that we observed a relationship between cluster preservation and memory performance but did not see a relationship of single voxel autocorrelation and memory performance, it is possible that examining the neural timescales alone is not sufficient to explain differences in clinical memory performance, and that the spatial location is a necessary component. For example, another study that characterized location-dependent patterns of functional connectivity throughout the hippocampus in healthy controls found a continuous gradient of functional connectivity along the long axis and that individual differences in the hippocampal gradient was related to performance on a recollection task (Przeździk et al., 2019). Those results converge with the current findings and suggest that examining the temporal component of voxels (i.e., single voxel autocorrelation) might provide limited information about the functional integrity of the signal and that combining it with spatial information (i.e., cluster preservation) results in a measure that shows greater sensitivity to variations in memory performance.

Consistent with findings from our previous work (Bouffard, Golestani et al., 2023; Coughlan et al., 2023), we found that data-driven clustering of the single voxel autocorrelation in TLE patients and controls resulted in three distinct clusters. Our clustering approach does not use a predefined number of clusters, and visual examinations of the cluster maps revealed that there were no differences in number of clusters between patients and controls, nor was there a difference between healthy and epileptogenic hemispheres. Other studies using data-driven clustering approaches of functional connectivity within the hippocampus have also found that TLE patients and controls have a similar number of clusters to one another and a similar number of clusters in both hemispheres (Barnett et al., 2019; Voets et al., 2014). This finding suggests that the total number of autocorrelation clusters is too coarse of a metric for differentiating between patients and controls or between healthy and epileptogenic hemispheres.

We had predicted that within patients, there would be differences in single voxel autocorrelation between the healthy and epileptogenic hemispheres. We did not find support for this prediction. The lack of difference in single voxel autocorrelation between hemispheres suggests that disruptions in intrinsic timescales of activity associated with left unilateral TLE extend into both hemispheres, which is consistent with prior work that found bilateral changes to intrinsic neural timescales in patients with TLE (Xie et al., 2023). Whereas we observed a significant difference in single voxel autocorrelation between LTLE and controls, there was no difference in single voxel autocorrelation between RTLE and controls. Null findings in patients with RTLE have appeared throughout the literature, suggesting that neural changes due to TLE manifest differently in left vs. right TLE. For example, cortico-hippocampal connectivity in LTLE was predictive of post-surgery outcomes, but not in RTLE (Audrain et al., 2023; Stasenko et al., 2023). Additionally, reductions in functional connectivity between the hippocampus and DMN regions are greater in LTLE compared with RTLE (Barnett et al., 2019; Pereira et al., 2010). Future work is needed to characterize the changes in temporal signaling in RTLE and how it is different from LTLE.

Mesial temporal sclerosis is associated with cell loss, which is especially prominent in the anterior hippocampus. We predicted that the structural changes due to MTS would affect the autocorrelation, particularly in the anterior-medial autocorrelation cluster. Contrary to our predictions we did not find significant effects of MTS and only found a marginal trend in patients with LTLE, where patients with MTS had slightly higher single voxel autocorrelation compared to those without. The functional gradients along the anterior-posterior and medial-lateral axes of the hippocampus have been shown to be coupled with the microstructure of the hippocampus (Vos de Wael et al., 2018), therefore there is strong reason to believe that structural changes due to MTS would affect the neural timescales of activity along the long axis of the hippocampus.

Our method did not detect differences in autocorrelation between MTS vs. no MTS, possibly due to the low resolution of the fMRI data or due to the small number of individuals with MTS in our patient sample (see limitations discussed below). Future work can be done to more explicitly test the relationship between changes in autocorrelation and the underlying microstructure of the hippocampus in individuals with and without MTS.

It is also plausible that single voxel autocorrelation is a measure of signal changes that cannot be fully explained by changes in structure. Recent work that examined intrinsic neural timescales in the hippocampus of patients with TLE found that when accounting for structural changes in their analyses (volume and cortical thinning) there was a reduced effect size, however patients still demonstrated reduced neural timescales compared to controls (Xie et al., 2023), suggesting that autocorrelation provides information about brain function that is distinct from brain structure. It is also possible that the impact of hippocampal structural changes due to MTS are not detectable during rest, and that the effects of structural changes become more apparent during hippocampally-dependent tasks, For example, TLE-related structural changes in the hippocampus mediated the reorganization of functional memory networks during episodic retrieval states, but not during semantic retrieval states (Cabalo et al., 2023). This suggests that the impact of hippocampal structural integrity on functional memory networks might be more robust during episodic memory tasks that recruit the hippocampus.

A limitation of the current study was the resolution of the resting state data, which might have been too low to be able to isolate and exclude voxels that had cell loss. We chose to resample the data (native resolution: 3.75 x 3.75 x 5 mm) to 2 mm isotropic so that we could align all individuals to a common MNI space, which was necessary for our clustering and cluster preservation analyses. Resampling, however, does not improve the native resolution of the data. The low-resolution data makes it difficult to differentiate between voxels with and without sclerosis. Further, we cannot account for partial volume effects, where a single voxel might contain healthy tissue as well as CSF where sclerosis has occurred. These effects, however, would likely amplify, rather than obscure, differences between the MTS and no MTS groups, therefore, partial volume effects are an unlikely explanation for our null findings. Future work combining high resolution 7T fMRI and advanced automated hippocampal segmentation is needed to precisely quantify change in single voxel autocorrelation and relate it to structural changes caused by MTS.

Another limitation is the small number of patients with MTS, leading to lower statistical power which potentially decreases our ability to find an effect of MTS. We attempted to address this by conducting the flipped brain analysis where we normalized both groups to the control group and then used a grouping variable of “epileptogenic hemisphere” and “healthy hemisphere” instead of left and right hemisphere (Liu et al., 2015), however we still did not observe significant effects of epileptogenic vs. healthy hemisphere on the average autocorrelation. It is possible that future studies with greater sample sizes might have more statistical power to investigate this effect.

These findings may be interpreted in the context of large-scale network disorganization in TLE. It is well-established that TLE disrupts connectivity patterns both within the hippocampus and across networks such as the default mode network (DMN) and medial temporal lobe memory systems (Bernhardt et al., 2013; Holmes et al., 2014; McCormick et al., 2013; Roger et al., 2020). There is evidence that the autocorrelation of individual voxels is associated with disruptions in network connectivity that is directly related to decreased memory performance in TLE (Nedic et al., 2015). Nedic et al. found a cluster of slow autocorrelation decay in the ipsilateral anterior hippocampus of TLE patients. In patients, this cluster had decreased connectivity with regions of the DMN compared to the same cluster in healthy controls, and the extent of connectivity decrease was related to worse memory performance on neuropsychological tests. Our finding that cluster-level spatial disorganization of autocorrelation patterns is related to memory performance could reflect or contribute to underlying instability in how the hippocampus participates in these broader functional networks. This disruption may manifest as both abnormal temporal dynamics locally and altered network-level communication globally – a possibility that warrants further research.

Importantly, we do not propose that single voxel autocorrelation should replace established structural or electrophysiological markers of hippocampal pathology in TLE which provide critical information about disease severity and seizure localization.

Furthermore, while other in-vivo biomarkers of functional integrity including hippocampal volume and formation, task-related activation and hippocampal-neocortical connectivity have also been demonstrated to correlate with mnemonic performance and to predict memory outcomes following surgery, they do not capture the alterations in spatially organized dynamics within the hippocampus. As such, integrating voxel-wise autocorrelation with these other modalities could improve precision in characterizing hippocampal dysfunction.

Future multimodal studies might combine high-resolution fMRI, quantitative structural imaging, and intracranial EEG to better understand the links between signal dynamics, network integrity, and memory outcomes in TLE. This approach could help clarify whether abnormal autocorrelation reflects persistent excitability, impaired inhibition, or other physiological mechanisms. Moreover, mapping how these disruptions evolve over time (e.g., longitudinally across disease progression) could position autocorrelation as a candidate biomarker for functional reserve or cognitive risk.

## 5. Conclusion

By applying a fine-grained analysis of the autocorrelation of individual voxels in patients with TLE we determined that single voxel autocorrelation and autocorrelation clustering are affected by TLE. We found that in cases of left TLE there was a bilateral increase in intrinsic timescales in the anterior-medial hippocampus. We also show evidence that changes in the spatial organization of the anterior-medial autocorrelation cluster was related to verbal and visuospatial memory. This work advances our understanding of the properties of intrinsic hippocampal timescales and highlights the importance of examining the spatial organization of neural timescales, particularly in the anterior-medial hippocampus.

## Data and Code Availability

Data and code available upon request.

## Author Contributions

N.R.B., S.A., M.M., and M.P.M. contributed to the conception of the study. N.R.B. analyzed the data, prepared figures, and drafted the initial manuscript. S.A. and A.G. contributed to data preprocessing and analysis. M.P.M. provided the data and resources needed for this experiment. All authors contributed to the revisions of the manuscript.

## Funding

This research was funded by the Canadian Institute of Health Research (CIHR Project Grant 148762, to M.P.M) and by EpLink—The Epilepsy Research Program of the Ontario Brain Institute (OBI), a non-profit corporation funded partially by the Ontario government (M.P.M). This research was also supported by the Canadian Natural Sciences Engineering Research Council (Discovery Grant, RGPIN-2020-05747, to M.D.B.), the James S. McDonnell Foundation (Scholar Award to M.D.B.), the Canada Research Chairs Program (M.D.B.), and a Max and Gianna Glassman Chair in Neuropsychology (M.D.B.).

## Declaration of Competing Interests

None of the authors has any conflict of interest to disclose.

## Notes

### Competing Interest Statement

The authors have declared no competing interest.

### Summary of Updates

This version of the manuscript has updated based on peer review. The Introduction and Discussion were edited, a demographics table was added, and two new supplemental analyses were added.

